# Impact of herbivore preference on the benefit of plant trait variability

**DOI:** 10.1101/670158

**Authors:** Tatjana Thiel, Sarah Gaschler, Karsten Mody, Nico Blüthgen, Barbara Drossel

**Affiliations:** Institut für Festkörperphysik, Technische Universität Darmstadt, Hochschulst. 6, 64289 Darmstadt, Germany; Institut für Festkörperphysik, Technische Universität Darmstadtz, Hochschulst. 6, 64289 Darmstadt, Germany; Ecological Networks, Technische Universität Darmstadt, Schnittspahnstraße 3, 64287 Darmstadt, Germany

**Keywords:** Intraspecific trait variability, Herbivore preference, Plantherbivore model, Jensen’s inequality

## Abstract

We explore the hypothesis that intraspecific trait variability can be *per se* beneficial for the plant when the curvature of the herbivore response to this trait is concave downwards. This hypothesis is based on a mathematical relation for non-linear averaging (Jensen’s inequality), leading to reduced herbivory when the trait distribution becomes broader. Our study introduces and investigates a model for plants and their insect herbivores that includes an unequal distribution of nutrient content between leaves. In contrast to earlier publications, we take into account the ability of herbivores to choose leaves, and the associated costs. By performing computer simulations and analytic calculations, we find that this herbivore preference can considerably alter the conclusion cited above. In particular, we demonstrate that herbivore populations that show preference for leaves on which they grow well can benefit from large nutrient level variability independently of the curvature of the herbivore response function, and despite the cost for preference.

## 1 Introduction

Populations in nature consist of individuals that typically differ in size, physiology, morphology, behavior, and resource utilization (Gibert and Brassil, 2014; Schreiber et al., 2011). This intraspecific trait variability emerges due to two mechanisms, namely, (i) genetic diversity or variability (Albert et al., 2011, 2010b; Hughes et al., 2008; Gibert and Brassil, 2014) and (ii) the plastic response to different environmental conditions (i.e. phenotypic plasticity) (Albert et al., 2011, 2010b; Whitham et al., 2003). Furthermore, intraspecific trait variability can be classified according to three organization levels, namely, (i) population-level variability (i.e. populations of a single species differ in traits), (ii) inter-individual variability (or between-individual variability), and (iii) intra-individual variability (or within-individual variability) (Albert et al., 2011, 2010b; Bolnick et al., 2002). Despite the overwhelming empirical evidence of intraspecific trait variability on all of these organization levels (Herrera, 2009; Siefert et al., 2015; Jung et al., 2010; Fridley and Grime, 2010), ecological theory has focused mostly on *interspecific* trait variability, not taking into account that individuals of a species also differ in their traits.

In recent years, however, an increasing number of empirical and theoretical studies has underlined the importance of considering intraspecific trait variability (Violle et al., 2012; Whitham et al., 2003; Bolnick et al., 2011; Gibert and Brassil, 2014; Schreiber et al., 2011; Okuyama, 2008; Anderson et al., 1997; Doebeli, 1996, 1997; Doebeli and de Jong, 1999; Jung et al., 2010; Booth and Grime, 2003; Hughes et al., 2008; Herrera, 2017). Empirical studies show for instance that intraspecific genetic diversity enhances population stability (Agashe, 2009), yields increased species diversity (Booth and Grime, 2003; Hughes et al., 2008), and modulates community and ecosystem dynamics (Raffard et al., 2018).

Several theoretical studies implemented intraspecific trait variability by using quantitative genetics models (Bulmer et al., 1980; Bolnick et al., 2011) for traits such as predation efficiency (Doebeli, 1997), competition strength (Doebeli, 1996; Doebeli and de Jong, 1999; Drossel and McKane, 1999), and host-parasite coupling (Doebeli, 1996). Others implemented explicit trait distributions in population dynamics equations to take into account intraspecific trait variability in encounter or attack rates (Okuyama, 2008; Gibert and Brassil, 2014), handling times (Gibert and Brassil, 2014; Okuyama, 2008), competition coefficients (Vellend, 2006), and movement patterns (Anderson et al., 1997). These theoretical studies found for instance that intraspecific trait variability can considerably change the nature of the dynamics (Doebeli, 1996; Doebeli and de Jong, 1999), affect the stability (Okuyama, 2008; Gibert and Brassil, 2014; Doebeli, 1997), and increase the diversity (Vellend, 2006) of the system.

One hypothesis why intraspecific trait variability can have such strong effects on ecological systems is that intraspecific variability *per se* affects ecological dynamics (Bolnick et al., 2011) due to non-linear averaging via Jensen’s inequality (Okuyama, 2008; Bolnick et al., 2011; Ruel and Ayres, 1999; Wetzel et al., 2016). This mathematical theorem states that a concave upwards function (i.e. increasing slope, positive curvature) applied on a mean of a set of points is less or equal to the mean applied on the concave upwards function of these points (Jensen, 1906). The opposite is true when considering a concave downwards function (i.e. decreasing slope, negative curvature). Hence, populations with the same mean trait but with different trait variances can have different mean interaction strengths (Bolnick et al., 2011). This altered interaction strength can have a crucial impact on the stability and diversity of ecological systems (McCann et al., 1998; Rall et al., 2008; Valdovinos et al., 2010; Kondoh, 2006; Heckmann et al., 2012; Vos et al., 2004; Thiel et al., 2018). Indeed, Jensen’s inequality (or Jensen’s effect) is cited in different ecological contexts, for example in the case of a nonlinear relationship between attack rates and body sizes (Bolnick et al., 2011) or nutrient concentration and chemostat population growth (Bolnick et al., 2011). Furthermore, Jensen’s inequality is used to explain why variance in temperature elevates poikilotherm metabolic rates (concave upwards function) (Ruel and Ayres, 1999), why variance in light regimes depresses primary production (concave downwards function) (Ruel and Ayres, 1999), and why variance in tissue quality and secondary metabolites affects herbivore response (Ruel and Ayres, 1999).

Several authors (Wetzel et al., 2016; Ruel and Ayres, 1999) refer to Jensen’s inequality to explain the large variability in plant traits observable in nature (Herrera, 2009; Siefert et al., 2015; Coleman et al., 1994; Albert et al., 2010a,b). Indeed, plant individuals are known to differ in traits (Coleman et al., 1994; Siefert et al., 2015; Herrera, 2009; Albert et al., 2010b) such as height (Jung et al., 2010; Siefert et al., 2015; Albert et al., 2010a,b), leaf morphology (e.g. leaf area and thickness) (Siefert et al., 2015; Jung et al., 2010; Coleman et al., 1994; Albert et al., 2010b,a) and leaf chemicals, as for example leaf nitrogen and phosphorus concentration (Siefert et al., 2015; Albert et al., 2010a,b) or secondary metabolites (Ohmart et al., 1985; Ali and Agrawal, 2012; Hartmann, 1996; Moore et al., 2014). A large degree of intraspecific trait variability in plants is found on all organization levels, i.e., the population, inter-, and intraindividual level (Siefert et al., 2015; Herrera, 2009).

Intraspecific trait variability in plants can thus have considerable effects on the community and the ecosystem in which the plant lives (Jung et al., 2010; Whitlock et al., 2007). For instance on inter-individual level, there is evidence to suggest that intraspecific genotypic diversity causes increased plant productivity (Kotowska et al., 2010), herbivore performance or richness (Kotowska et al., 2010; Crutsinger et al., 2006), and species richness of higher trophic levels (Crutsinger et al., 2006). Furthermore, plant genotypic diversity can act as a barrier to invasive species (Crutsinger et al., 2008) or provide disease suppression (Zhu et al., 2000; Tooker and Frank, 2012). Nevertheless, modern agroecosystems are dominated by homogeneous monocultures. The lack of crop genetic diversity has considerable consequences for the ecosystem (Wetzel et al., 2016), as for instance increased herbivory (Tooker and Frank, 2012; Peacock and Herrick, 2000), decreased arthropod richness (Tooker and Frank, 2012; Crutsinger et al., 2006; Johnson et al., 2006), decreased plant fitness (Johnson et al., 2006; Tooker and Frank, 2012), increased pest and pathogen pressure (Tooker and Frank, 2012; Esquinas-Alcázar, 2005), and higher vulnerability to abrupt climate changes (Esquinas-Alcázar, 2005).

On the intra-individual level, however, less is known about the impact of trait variability. Wetzel et al. (2016) used Jensen’s inequality, which is in principle applicable on all organization levels of intraspecific trait variability, to argue that plant trait variability is *per se* beneficial for the plant when herbivore response is a concave downwards function of this trait. In this case, Jensen’s inequality states that the mean herbivore response is smaller when the trait values of plant leaves vary around a mean than when all leaves have the mean trait value. In a meta-study, Wetzel et al. (2016) indeed found that herbivore response is a concave downwards function of the leaf nutrient level. However, there are also several counterexamples (Ruel and Ayres, 1999). For instance, linear (Ayres et al., 1987) or complex herbivore response functions having both concave upwards and concave downwards regions (Clancy, 1992) were found in dependence of the nutrient level. A reason for these divergent results may be that the curvature of the herbivore response function depends on the considered nutrient, herbivore (Ali and Agrawal, 2012), nutrient level range (Clancy, 1992; Miles et al., 1982; Ohmart et al., 1985), and the age of the herbivore individuals (Scriber and Slansky Jr, 1981; Ohmart et al., 1985; Montgomery, 1982; Zalucki et al., 2002).

A fact not considered in the studies mentioned above is that herbivores are able to adapt to changes in their environment, such as the plant nutrient level distribution, via their behavior. For instance, herbivore preference is one important way to respond to trait distribution in the resource (Via, 1986; Herrera, 2009). It can appear in different forms: Herbivores can have *feeding* preference for leaves, on which they perform best. This may imply to feed on a mixture of plants and requires mobility to reach appropriate leaves (Mody et al., 2007; Lubchenco, 1978). Additionally, *oviposition* preference for leaves with certain traits is regularly found in nature (Via, 1986; Herrera, 2009; Tabashnik et al., 1981; Travers-Martin and Müller, 2008; Despres et al., 2007; Rausher, 1979). Here, several studies support the so-called preference-performance hypothesis (or “mother-knows-best hypothesis”) which states that herbivores choose egg-laying sites where their offspring perform best (Soto et al., 2012; Tilmon, 2008; Gripenberg et al., 2010), although there are several studies that found mismatches between herbivore preference and performance (Valladares and Lawton, 1991; Gripenberg et al., 2010; Hufnagel et al., 2017). Reasons for these mismatches may be that adults are not able to properly discriminate the leaf traits (e.g. due to associational effects (Barbosa et al., 2009)), try to reduce larvae competition (Wetzel and Thaler, 2018), or choose oviposition sites where they perform best that deviate from the best feeding sites for their offspring (Scheirs et al., 2000; Scheirs and De Bruyn, 2002). Furthermore, temporal variation of the plant traits or strategies to avoid predators (Björkman et al., 1997) may be responsible for these mismatches. The organization level of herbivore preference, i.e. whether the herbivores prefer leaves (i) of a certain plant population, (ii) of a certain plant individual, or (iii) with a certain trait within a single plant individual, is determined by the organization level of plant trait variability.

Up to now, theoretical models for adaptive feeding behavior focus on consumers (Valdovinos et al., 2010) that choose among several prey or resource species: In food web models, preference in form of adaptive foraging (Kondoh, 2006; Heckmann et al., 2012; Uchida et al., 2007; Feng et al., 2009) or adaptive prey switching (Fasham et al., 1990; Valdovinos et al., 2010) was shown to enhance the stability and species diversity of the food web. In a plant-herbivore model with feeding preference for the most edible plant species (Grover and Holt, 1998; Feng et al., 2009), it was found that the herbivores drive the plant with the less effective toxin to extinction (Feng et al., 2009). Adaptive herbivory thus reduces local species diversity (Feng et al., 2009). However, less is known about the impact of herbivore preference on an intraspecific (or even intra-individual) level, although several empirical studies find that preference occurs on this level (Mody et al., 2007; Gutbrodt et al., 2012; Rausher, 1979). In particular, it has not been explored so far how herbivore preference affects the impact of intraspecific plant trait variability on herbivore fitness.

In this paper, we want to fill this gap. We propose a plant-herbivore model that includes plant nutrient level variability and herbivore preference and is valid for intra-individual and inter-individual nutrient level variability as well as for feeding and oviposition preference. Based on the study of Wetzel et al. (2016), we investigate whether plant nutrient level variability is *per se* beneficial or disadvantageous for a herbivore population depending on the curvature of the herbivore response function. In particular, we explore how herbivore preference affects this result. We couple the extent of the preference with corresponding costs for finding appropriate leaves (Tilmon, 2008).

In order to distill the effect of intraspecific trait variability *per se*, we focus on one trait, namely the nutrient level in the plant leaves, and neglect possible correlated variations in secondary metabolites. For the same reason, we only model the herbivore population and neglect higher trophic levels. Furthermore, we assume that the plant population is sufficiently large that it can be considered as constant over the time span covered by the model. Furthermore, we consider response functions with different curvatures in our study in order to take for instance different types of nutrients, nutrient level ranges (Miles et al., 1982), and ages of the herbivore individuals (Scriber and Slansky Jr, 1981; Ohmart et al., 1985; Montgomery, 1982) into account.

We show that herbivore preference crucially affects the predictions of (Wetzel et al., 2016) that are based on Jensen’s inequality. More precisely, we find that when a herbivore population has a strong preference, it benefits from large plant trait variability, irrespective of the curvature of the response function. In addition to computer simulations, we show this also analytically. Furthermore, we evaluate the optimal herbivore preference for a given plant nutrient level variability and different herbivore response functions. Here, we show that the curvature of the response function determines the strength of optimal herbivore preference.

## 2 Model

We consider insect herbivores feeding on a plant population whose leaves show a distribution *p*(*n*) of the nutrient level *n*. More precisely, we assume that the nutrient level *n* describes the food quality from the herbivore’s point of view and thus the quality of nutrient composition. We therefore always consider monotonically increasing herbivore response functions. We further assume that there is always enough consumable plant material such that intraspecific competition for food is negligible.

### 2.1 Herbivore fitness

The central quantity to be evaluated is herbivore population fitness, since it is a direct indicator of herbivore population growth. The mean fitness 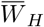 of a herbivore population is defined as the mean number of offspring per herbivore individual reaching reproductive age. Denoting the distribution of herbivore individuals on leaves with nutrient level *n* as *Φ*(*n*) and the fitness of a herbivore individual feeding on a leaf with nutrient level *n* as *W*_*H*_ (*n*), the mean population fitness can be expressed as

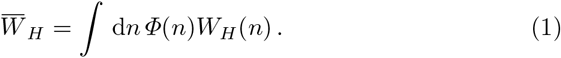

The fitness *W*_*H*_ (*n*) of a herbivore individual feeding on a leaf with nutrient *n* depends on the growth of the herbivore on this leaf, which we will express in terms of a performance function *f* (*n*). We define the performance function *f* (*n*) as the weight gain of a herbivore individual feeding on a leaf with nutrient level *n* from hatching to pupation. The different types of performance functions used in our study will be specified further below. If we assume that the number of offspring that reach reproductive age is proportional to this weight gain, the fitness *W*_*H*_ (*n*) of a herbivore individual feeding on a leaf with nutrient level *n* is given by

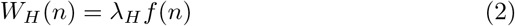

with *λ_H_* being the number of offspring per unit of weight gain.

### 2.2 Distribution of herbivore individuals on leaves

The distribution *Φ*(*n*) of herbivore individuals on leaves with nutrient level *n* depends on the one hand on the nutrient distribution *p*(*n*), and on the other hand on a preference function *Φ*_*p*_(*n*) that quantifies the extent of preference for leaves with nutrient level *n*. Since we assume that intraspecific competition for food is negligible, the distribution is obtained by multiplying the preference function with the leaf abundance and normalizing the result, leading to

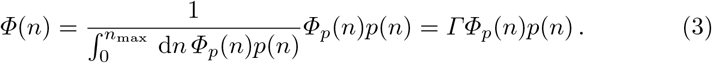

We use Gaussian distributions in order to describe both the nutrient distribution *p*(*n*) and the preference function *Φ*_*p*_(*n*). More details are given further below.

### 2.3 Nutrient distribution

For the nutrient distribution *p*(*n*) among leaves, we assume a Gaussian distribution with a mean in the middle of the considered nutrient level interval *n* ∊ [0, *n*_max_], i.e. at 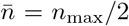. The variance *V_S_* of this distribution determines the degree of heterogeneity of the nutrient distribution. Depending on the herbivore preference (s. Section 2.4 for details) and the herbivore performance function (s. Section 2.5 for details), it may be favorable for the plant population to have a broad or a narrow nutrient distribution. This distribution thus represents a strategy of the plant (Wetzel et al., 2016). We introduce the strategy parameter

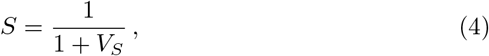

such that *S* = 0 represents a uniform distribution over the considered nutrient level interval and *S* = 1 a delta distribution, meaning that all leaves of all plant individuals have the nutrient level 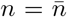. Fig. 1(b) shows the nutrient distribution *p*(*n*) for different plant strategies *S*.

**Fig. 1.**
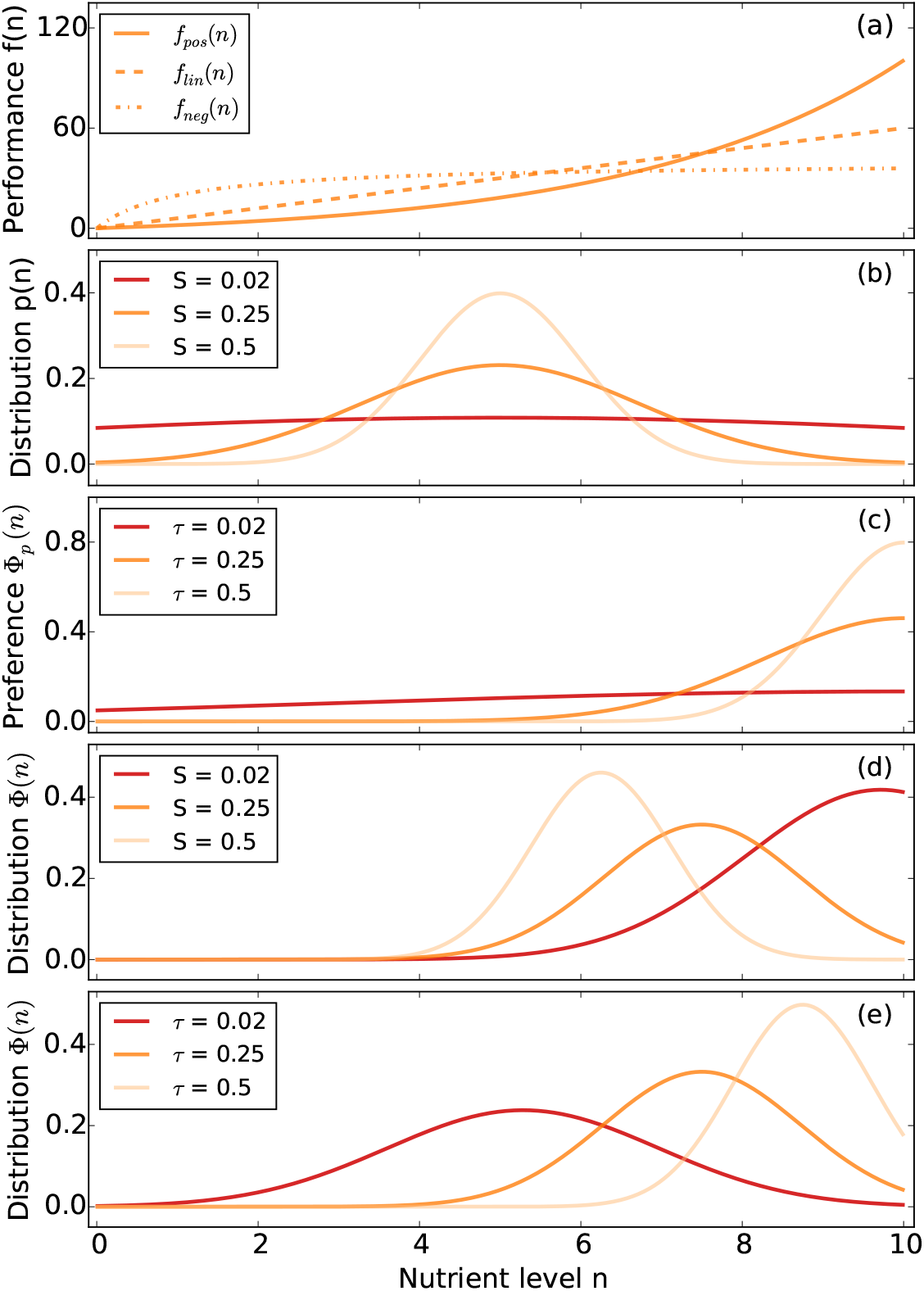
Graphical illustration of the different functions occurring in our model. *(a)* The three types of herbivore performance functions considered in our study: *f*_pos_(*n*) = 0.12*n*^3^, *f*_lin_(*n*) = 6*n*, 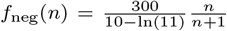. *(b)* Change of the nutrient distribution *p*(*n*) with plant strategy parameter *S* (low *S* means high nutrient level variability, high *S* low variability; cp. Eq.(4)). *(c)* Preference function *Φ*_*p*_(*n*) for different strengths of preference *τ* (cp. Eq.(5)). *(d)* Distribution *Φ*(*n*) of herbivore individuals on leaves with nutrient level *n* (cp. Eq.(3)) for different plant strategies *S* (cp. Eq.(4)) and a herbivore preference *τ* = 0.25 (cp. Eq.(5)). This is the (normalized) product of the orange curve in (c) with the three different curves in (b). *(e)* Distribution *Φ*(*n*) of herbivore individuals on leaves with nutrient level *n* (cp. Eq.(3)) for different herbivore preferences *τ* (cp. Eq.(5)) and a plant strategy parameter *S* = 0.25 (cp. Eq.(4)). This is the (normalized) product of the orange curve in (b) with the three different curves in (c).

### 2.4 Preference function

The preference function *Φ*_*p*_(*n*) can be interpreted as the probability that an adult herbivore lays eggs on a leaf with a nutrient level *n* when encountering it. We model this function via a Gaussian distribution with its mean at the performance maximum of the herbivore population and a variance *V*_*p*_ that is smaller when the preference is stronger. Hence, herbivores prefer leaves on which they or their offspring perform well as observed in several studies (Via, 1986; Herrera, 2009; Tabashnik et al., 1981; Travers-Martin and Müller, 2008; Despres et al., 2007; Rausher, 1979). Note that our results do not qualitatively depend on the position of the preference mean as long as herbivores prefer highquality leaves. We define the preference parameter

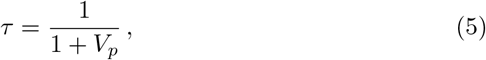

such that *τ* = 0 stands for no, and *τ* = 1 for full preference. In the latter case, preference function is a delta distribution which describes the unrealistic extreme case that only those leaves are used for oviposition on which the herbivore population reaches its performance maximum. Fig. 1(c) shows the preference function *Φ*_*p*_(*n*) for different preferences *τ*. Instead of oviposition preference, the preference function can also indicate a feeding preference. In this case, the preference function *Φ*_*p*_(*n*) describes the probability that a herbivore feeds on a leaf with nutrient level *n* when encountering it. Note that the preference function is a population average, such that diverging preferences of herbivore individuals or an incapability to discriminate leaf traits properly lead to a low population preference *τ*. Furthermore, less herbivore individuals feed on high-quality leaves when these become rare (i.e. with increasing plant strategy parameter *S*), even when the value of *τ* (i.e., the degree of preference) is not changed, since less herbivore individuals encounter these high-quality leaves (cp. Fig. 1(d)).

Preference comes with a cost for finding appropriate leaves. We take this cost into account in form of a mass loss of the herbivore. Since we want to explore the effect of this cost, we describe the relative mass loss by a function that allows us to interpolate between 0 and 1 in different ways by changing the parameters of this function. In this way, we can make sure that unrealistic extreme cases of the preference function, such as a delta distribution, do not lead to survival of the herbivore population. We thus define the relative mass loss due to preference as

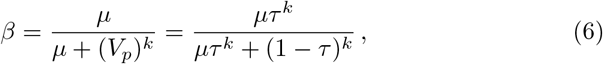

where larger *μ* means that the cost of preference is larger, and large *k* means that the costs are mainly incurred when preference is large. Fig. 8 in the Appendix shows the proportion of remaining mass considering preference, 1 − *β*, as a function of preference *τ* for different values for *μ* and *k*.

Including this cost changes the expression Eq.(1) for the mean fitness of the population to

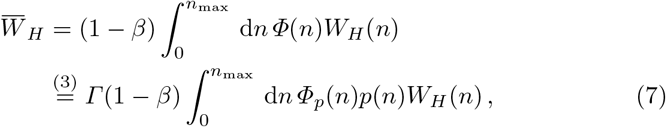

where *Γ* normalizes again the distribution *Φ*(*n*) of herbivore individuals on leaves with nutrient *n* to 1. Figs. 1(d), (e) show the distribution *Φ*(*n*) of herbivore individuals on leaves with nutrient *n* for varying plant strategy parameter *S* and varying preference *τ*, respectively.

### 2.5 Performance function

Wetzel et al. (2016) argued that the curvature of the herbivore performance function determines via non-linear averaging (Jensen’s inequality) whether a herbivore benefits or suffers from plant trait variability. In order to test this idea, we will perform our computer simulations using three performance functions with different curvature, which are shown in Fig. 1(a). We chose them such that the mean performance is the same in the three cases in order to ensure that the resulting mean fitness values are of the same order of magnitude.

All these functions are plausible, and they apply to different types of nutrients and to different ranges of their concentration: The concave upwards function (continuous line) is appropriate when the nutrient limits growth. The linear function (dashed line) is suitable when growth is additionally dampened for instance by a conversion process that requires energy. Eventually, herbivore growth should saturate with increasing nutrient level, leading to the concave downwards form (dot-dashed line). This scenario applies when considering a nutrient that is important for growth, but typically does not limit growth, or when the considered nutrient level interval is so large that absolute limits to growth become visible.

Under the assumption that herbivore performance functions increase monotonically, the functions shown in Fig. 1(a) represent all possibilities that differ qualitatively in their curvature. We do not consider s-shaped functions that include both concave upwards and concave downwards regions because we want to focus on the effect of the curvature of the performance function on herbivore fitness in order to test Jensen’s inequality. Performance functions that decrease again for large nutrient concentrations can occur when excess nutrients lead to negative effects. In fact, empirical studies often found performance functions that have a form like a concave downwards parabola with a maximum at an intermediate nutrient level, for nutrients such as nitrogen (Zehnder and Hunter, 2009; Joern and Behmer, 1998; Fischer and Fiedler, 2000; Joern and Behmer, 1997) or phosphorus (Boersma and Elser, 2006). In literature, three reasons have been proposed to explain this observation (Tao et al., 2014), namely (i) correlated changes in other physical/chemical properties (Boersma and Elser, 2006; Raubenheimer et al., 2009), (ii) increased metabolic costs for excreting or storing excess nutrients, potentially being the reason why it is observed that (iii) species adapt their total intake to avoid strong excess of one nutrient despite resulting limitation of another essential nutrient (Raubenheimer et al., 2005; Lee et al., 2004).

As we assume that the nutrient level *n* describes the food quality from the herbivore’s point of view, these effects are not relevant for our model. To allow a comparison with unimodal responses of herbivores to particular nutrient levels in plants (e.g. (Wetzel et al., 2016)), we consider in the Appendix nevertheless performance functions that have their maximum at an intermediate nutrient level (s. Section A.1).

### 2.6 Choice of parameter values

We chose the nutrient level range to be *n* ∈ [0, 10], such that the mean nutrient level is 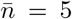. By choosing appropriate units for the nutrient level, every nutrient level interval can be mapped onto this one. As we want to investigate whether nutrient level variability is *per se* beneficial or disadvantageous for the herbivore population, we keep the mean nutrient level constant and just alter the plant strategy parameter *S*, i.e. the variance of the nutrient level distribution.

For the cost of preference, Eq.(6), we choose the parameters *μ* = 1 and *k* = 2, such that the costs for a moderate level of preference remain moderate, but become considerable for high preference. In order to find an appropriate value for the number of offspring per unit of growth, *λ*_*H*_, we choose the forest tent caterpillars (*Malacosoma disstria*) as model species. *Malacosoma disstria* has a typical mass gain until pupation of 300 mg and produces around 300 eggs with a survival rate of 1/100 resulting in a number of offspring reaching the reproductive age per growth unit of 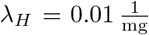 (Hemming and Lindroth, 1999). Furthermore, we normalize the mean performance of all functions to 300 mg, i.e. 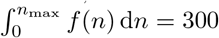. Note that these parameter values only have a quantitative impact on the fitness landscape of the herbivore population.

## 3 Results

We divide our investigation into two parts. In a first step, we analyze the herbivore fitness (cp. Eq.(7)) in dependence of the plant strategy parameter *S* (cp. Eq.(4)) and the shape of the performance function without herbivore preference, i.e. for *τ* = 0 (cp. Eq.(5)). These results can be compared to previous studies (Wetzel et al., 2016; Ruel and Ayres, 1999; Bolnick et al., 2011) and will act as a reference for the second part of our investigations. In the second step, we investigate the effect of herbivore preference *τ* > 0 on these results, and we will show that herbivore preference has a crucial impact on the conclusions and assumptions in (Wetzel et al., 2016; Ruel and Ayres, 1999; Bolnick et al., 2011).

### 3.1 Herbivore fitness in dependence of plant strategy in absence of herbivore preference

We first consider the situation that the herbivore population shows no preference, i.e., *τ* = 0 and *β* = 0. Hence, the preference function *Φ*_*p*_(*n*) is a uniform distribution such that the distribution of herbivore individuals on leaves with nutrient level *n* is *Φ*(*n*) = *Γp*(*n*) (cp. Eq.(3)) and the expression for the mean fitness (cp. Eq.(7) and Eq.(2)) simplifies to

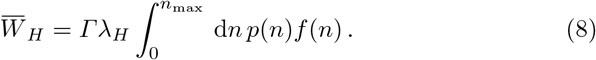

Fig. 2 shows the herbivore fitness (cp. Eq.(8)) in response to the plant strategy parameter *S* (i.e. the width of the nutrient distribution; cp. Eq.(4)) using the three different performance functions presented in Section 2.5 (cp. Fig. 1(a)).

**Fig. 2.**
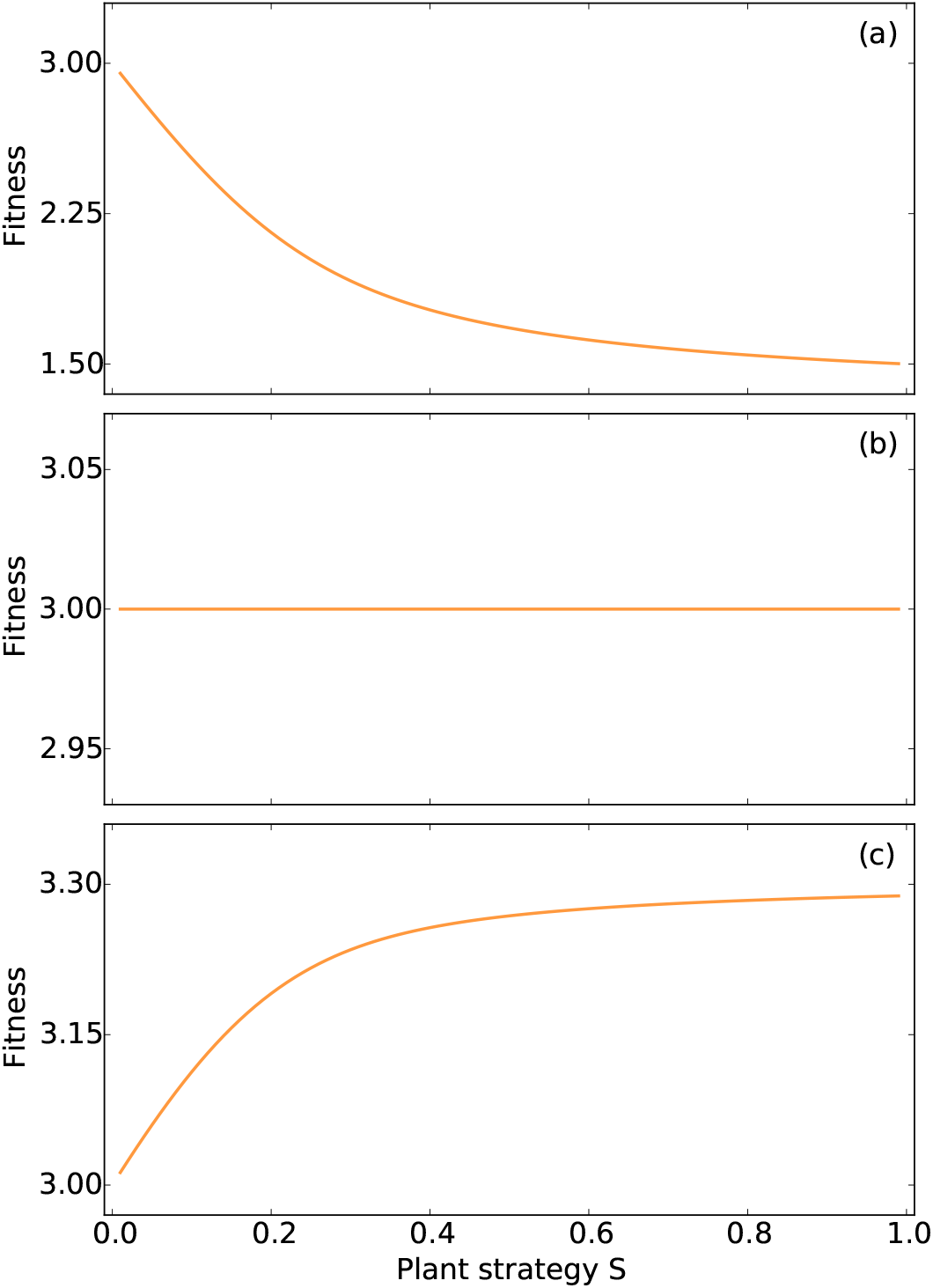
Mean herbivore population fitness (i.e. mean number of offspring per herbivore individual reaching reproductive age; cp. Eq.(8)) in dependence of the plant strategy parameter *S* (low *S* means high nutrient level variability, high *S* low variability; cp. Eq.(4)) for the case that the herbivore shows no preference (i.e. *τ* = 0) for the three different types of performance functions, *(a)* the concave upwards performance function *f*_pos_(*n*); *(b)* the linear performance function *f*_lin_(*n*); *(c)* the concave downwards performance function *f*_neg_(*n*) (cp. Fig. 1(a)).

The curvature of the performance function determines whether the herbivore population benefits or suffers from high nutrient level variability (i.e. small *S*) as expected: in the case of a concave upwards performance function *f*_pos_ the herbivore population is fitter when nutrient level variability is larger while the opposite is true in the case of a concave downwards performance function *f*_neg_. For a linear performance function *f*_lin_, the plant strategy parameter *S* has no influence on herbivore population fitness.

These results can be understood using Jensen’s inequality (Jensen, 1906), which is based on non-linear averaging. This mathematical theorem states that a concave upwards function of the mean value of a set of points *x*_*i*_ is less or equal to the mean value of the concave upwards function of these points, i.e. 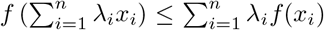 with a concave upwards function *f* and 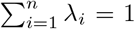 with *λ*_*i*_ ≥ 0 ∀*i* ∊ [1, *n*]. The opposite is true for a concave downwards function, and for a lin ar function *f* both values are the same. Consequently, we can deduce that 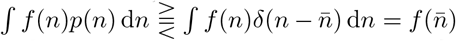, depending on the curvature of *f*, whenever *S* < 1, which means that the nutrient distribution has a nonzero width (cp. Eq.(4)). The difference between the two sides of the inequality becomes larger when the curvature of *f* is larger in the relevant range of *n* values, as can be seen in Fig. 2.

For further understanding, we reconstruct these results analytically. Additionally, the calculation will act as a basis for the following part, where we investigate the impact of preference on these results. We calculate the mean fitness (cp. Eq.(8)) as a function of the plant strategy parameter *S* (cp. Eq.(4)) when the herbivore has no preference (i.e. *τ* = 0) and expand the result for small variations of the nutrient level *n* around the mean nutrient level 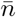. In order to do this, we approximate the performance functions by a polynomial in *n*, the curvature of which depends on a parameter *c*, i.e.

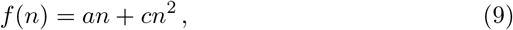

with the curvature parameter *c* and *a* ≥ 0. The performance function is concave upwards when *c* > 0, linear when *c* = 0, and concave downwards when *c* < 0. Both performance functions *f*_pos_ and *f*_neg_ (cp. Fig. 1(a)) can be transformed to the performance function defined above with *c* > 0 and *c* < 0, respectively, by using a Taylor expansion around the mean nutrient level 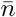, appropriately choosing the parameter *a*, and neglecting constant terms since they have no qualitative impact on the fitness landscape. Starting with Eq.(8), we find

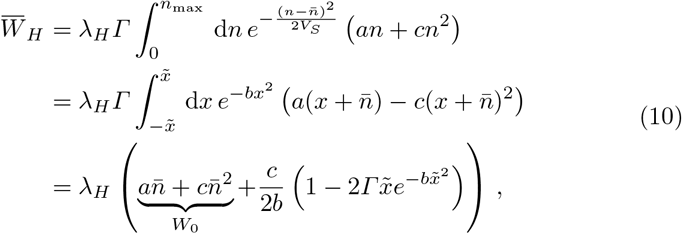

where we substituted 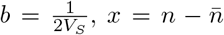 and used integration by parts to solve the integral. Hence, for small variations around the mean nutrient level, i.e. around 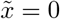, we find with 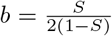 (cp. Eq.(4))

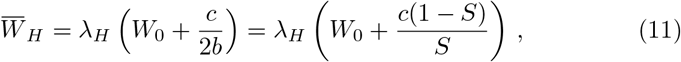

such that the sign of the curvature parameter *c* determines whether the mean fitness increases or decreases with increasing nutrient level variability, i.e. as a function of *b* or *S*. This also shows why mean herbivore fitness does not depend on the nutrient level variability when considering a linear performance function *f*_lin_ (i.e. *c* = 0). Note that this calculation also includes performance functions that do not increase monotonically, as the performance function for negative *c* becomes a concave downwards parabola over the considered range of nutrient concentrations when |*c*| is sufficiently large.

### 3.2 Herbivore fitness in response to herbivore preference and plant strategy

As a second step, we now take herbivore preference into account. Fig. 3 shows the mean herbivore population fitness 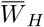 (cp. Eq.(1)) displayed in color code as a function of the herbivore preference *τ* (cp. Eq.(5)) and the plant strategy parameter *S* (cp. Eq.(4)) using (a) *f*_pos_, (b) *f*_lin_, and (c) *f*_neg_ as performance function (cp. Fig. 1(a)).

**Fig. 3.**
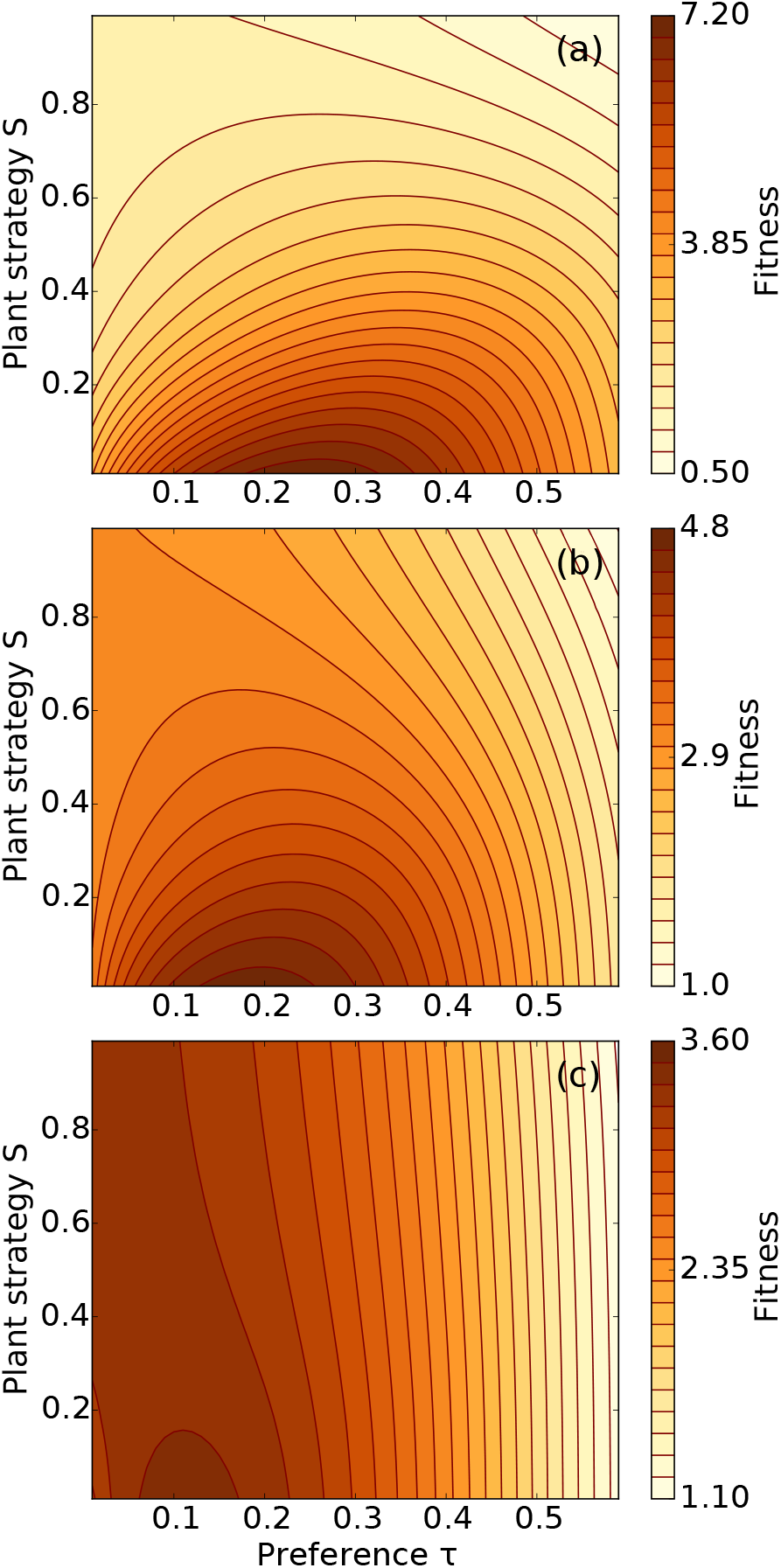
Herbivore fitness (i.e. mean number of offspring per herbivore individual reaching reproductive age; cp. Eq.(7)) displayed in color code in dependence of the plant strategy parameter *S* (low *S* means high nutrient level variability, high *S* low variability; cp. Eq.(4)) and herbivore preference *τ* (cp. Eq.(5)) considering *(a)* the concave upwards performance function *f*_pos_(*n*); *(b)* the linear performance function *f*_lin_(*n*); *(c)* the concave downwards performance function *f*_neg_(*n*) (cp. Fig. 1(a)).

When herbivores have a preference for leaves with more nutrients, the curvature of the performance function is not the only factor determining whether the herbivore population benefits or suffers from large plant nutrient level variability. The mean herbivore fitness decreases with increasing plant strategy parameter *S* (i.e. a narrower nutrient distribution; cp. Eq.(4)) for all possible preferences *τ* ∈ (0, 1) when the performance function is concave upwards *f*_pos_ (cp. Fig. 3(a)), as it can be seen from the change from darker to a lighter color with increasing *S*. However, there is no such uniform trend for a linear or concave downwards performance function. More precisely, the mean herbivore population fitness decreases with increasing plant strategy parameter *S* as soon as the herbivore population has a nonzero preference *τ* when the performance function is linear *f*_lin_ (cp. Fig. 3(b) and Fig. 2(b)) as illustrated by color changes from darker to a lighter color with increasing *S*; with a concave downwards performance function *f*_neg_, this transition point is reached when *τ* ⪆ 0.05 (cp. Fig. 3(c)) with our choice of parameters. This means that the fitness varies less under changes of the nutrient level variability when approaching the transition point and is (nearly) independent of *S* at the transition point. Note that the total amount of nutrients being available for the herbivore population is kept constant in our investigation and that this effect arises just due to a redistribution of nutrients between the leaves.

The fitness change with the plant strategy *S* is largest for the concave upwards performance function followed by the linear and the concave downwards performance function. This is the case since the concave upwards performance function changes most in the relevant range, i.e. for *n* ≥ 5 (cp. Fig. 1(d), (e)). For the same reason the fitness varies less when the nutrient distribution becomes narrower (i.e. larger *S*).

In Section A.1 in the Appendix, we repeated this investigation with a performance function having its maximum at an intermediate nutrient level. We found that our results do not change qualitatively. In the special case that the mean nutrient level coincides with the maximum of the performance function, an increase in nutrient variability always leads to decreased herbivore fitness, no matter how low the cost of preference is.

#### 3.2.1 Analytic calculations with preference

In the previous section, we found that herbivore populations benefit from large nutrient level variability irrespective of the curvature of the performance function when having considerable preference for leaves with high nutrient level. In order to understand this result and to demonstrate that this finding is generic, we calculate the mean fitness 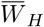 (cp. Eq.(1)) under the assumption that the nutrient level distribution is narrow. In this case, the preference function *Φ*_*p*_(*n*) can be Taylor expanded, and for a small preference value *τ* we have

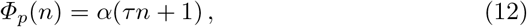

where *α* is a scaling factor. As in the previous section, we use the performance function *f* (*n*) = *an* + *cn*^2^, with the curvature parameter *c*. We find

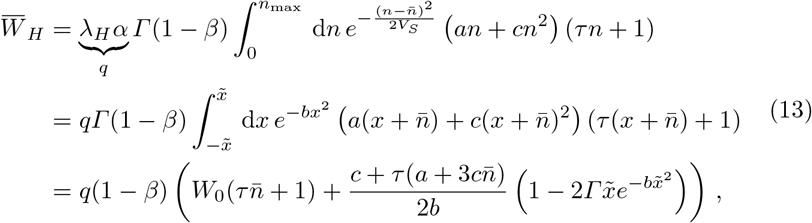

where we again substituted 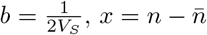 and used integration by parts to solve the integral. The mass loss due to preference *β* is proportional to the preference *τ*, i.e. *β* = *γτ*. Hence, for small variations around the mean nutrient level, i.e. around 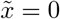, for small curvature parameter *c*, and by keeping in mind that we consider small preference values *τ*, we find with 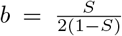(cp. Eq.(4))

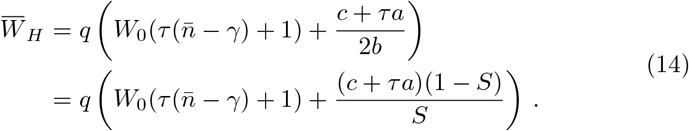

Consequently, as soon as the herbivore population exhibits some preference (*τ* > 0), the mean fitness 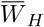 increases with increasing nutrient level variability (i.e. decreasing *S* or *b*) when the performance function is linear (i.e. *c* = 0). This means that the herbivore population benefits from large nutrient level variability. Furthermore, when we consider a concave downwards performance function *f*_neg_ (i.e. *c* < 0), the mean fitness increases with increasing variability (i.e. decreasing *S* or *b*) when |*c*| < *τa*. Hence, when |*c*| = *τa*, herbivore fitness is independent of the plant strategy parameter *S*. This corresponds to the observations from computer simulations shown in the previous section.

#### 3.2.2 Optimal herbivore preference for a given plant strategy

Herbivore preference is a flexible strategy that can be adapted to environmental conditions. In an evolutionary process, the herbivore population would evolve to the preference strategy that maximizes the mean population fitness. In Fig. 4, we plot this optimal preference *τ* in dependence of the plant strategy parameter *S* (cp. Eq.(4)) for the concave upwards *f*_pos_(*n*) (diamonds), linear *f*_lin_(*n*) (circles), and concave downwards *f*_neg_(*n*) (triangles) performance function shown in Fig. 1(a). The color shades in the markers display the mean fitness of the herbivore population under the particular circumstances.

**Fig. 4.**
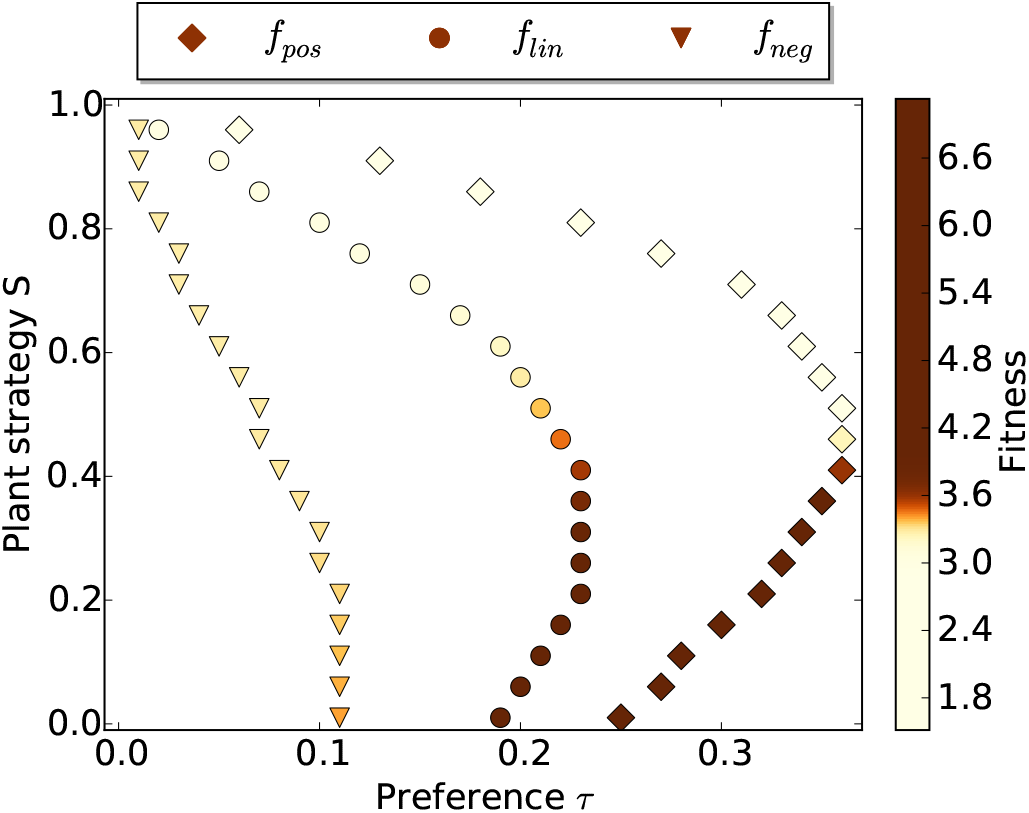
Herbivore preference *τ* that maximizes herbivore fitness for a given plant strategy parameter *S* (low *S* means high nutrient level variability, high *S* low variability; cp. Eq.(4)), for the three different performance functions *f* (*n*): the concave upwards performance function *f*_pos_(*n*), the linear performance function *f*_lin_(*n*), and the concave downwards performance function *f*_neg_(*n*) (cp. Fig. 1(a)). The color shades in the markers display the mean fitness of the herbivore population.

When all nutrient levels occur equally often in the plant leaves (i.e. *S* = 0), herbivore fitness is maximal at an intermediate preference value *τ*, since a moderate degree of preference already leads to a large gain in nutrients, and higher preference incurs higher energetic costs for searching high-nutrient leaves (cp. Fig. 4 and Fig. 3).

The fitness that is reached with optimal herbivore preference decreases with increasing plant strategy parameter *S* (cp. Eq.(4)) as can be seen from the color changes from darker to lighter color (s. Fig. 4). This is the case since preference has the largest impact on the distribution *Φ*(*n*) of herbivores on leaves with nutrient level *n* when the nutrients are equally distributed (cp. Fig. 1(d), (e)). The change in mean fitness as *S* increases from 0 to 1 is largest for the concave upwards performance function, *f*_pos_, as can be seen from the color changes from very dark to very light color, and is only moderate when the performance function is concave downwards, *f*_neg_, suggesting that in this case the food intake of a herbivore population depends less on the plant strategy.

For each value of the plant strategy parameter *S*, the optimal preference value *τ* is largest for the concave upwards performance function *f*_pos_(*n*) followed by the linear performance function *f*_lin_(*n*), and is smallest for the concave downwards performance function *f*_neg_(*n*) (cp. Fig. 4 and Fig. 3). This is plausible by looking at the shape of the performance functions (cp. Fig. 1(a)): In the case of the concave upwards performance function *f*_pos_(*n*), herbivore performance (and therefore herbivore growth) increases considerably when consuming leaves with higher nutrient concentration, since the performance function becomes steeper with increasing *n*. In contrast, with a concave downwards performance function, *f*_neg_(*n*), the consumption of leaves with higher nutrient concentration increases performance much less, since the performance function becomes flatter with increasing *n*.

When the nutrient distribution is very narrow (i.e. high *S*), the optimum preference value is small for all three types of performance function, since high-nutrient leaves are rare such that the costs of finding them exceeds the benefit.

In the Appendix (s. Section A.3), we show that these results are robust under changes of the parameters that shape the mass loss due to preference *μ* and *k* (cp. Eq.(6)). We also show that the optimum preference value increases when the curvature of the performance function becomes larger (s. Section A.2).

## 4 Discussion

In the present paper, we proposed a plant-herbivore model that includes herbivore preference and plant nutrient level variability, and we investigated the impact of these features on herbivore fitness depending on the curvature of the herbivore performance function. Our model and our results are valid for intra- and inter-individual nutrient level variability as well as for oviposition preference and feeding preference in the larval stage. The trait variability in our model describes spatial variation of the nutrient concentration, for instance due to diverging environmental conditions or development stages of the leaves. Thereby, we distilled the effect of nutrient level variability by considering a constant mean nutrient level. Note that changing mean nutrient concentrations may have additional effects (Wetzel and Thaler, 2018).

As we considered the nutrient quality from the herbivore’s point of view, our study focuses on performance functions with a positive slope. However, we made sure that the results obtained for concave downwards performance functions apply also to the case where the performance function has the shape of a concave downwards parabola with a maximum at intermediate nutrient levels (when herbivores have no preference or when the performance maximum is not close to the mean nutrient level).

In the case of no herbivore preference (*τ* = 0), we found that the herbivore population suffers from plant nutrient level variability when the herbivore performance function is concave downwards (i.e. decreasing slope, negative curvature), but benefits in the case of a concave upwards performance function (i.e. increasing slope, positive curvature) (cp. Fig. 1(a)). Indeed, this can be shown by a simple analytic calculation. So far, our findings agree with other authors who argue, based on Jensen’s inequality, that the curvature of the performance function determines via non-linear averaging whether the herbivore population benefits or suffers from plant trait variability (Wetzel et al., 2016; Bolnick et al., 2011; Ruel and Ayres, 1999).

Wetzel et al. (2016) performed a meta-study in order to obtain empirical herbivore performance functions. They found that herbivore performance is on average a concave downwards function of the plant nutrient level. The authors infer that this may explain the large nutrient level variability seen in plants (Herrera, 2009; Siefert et al., 2015), since plants can reduce the fitness of their herbivores by increasing nutrient level variability.

However, other studies found linear performance functions (Ayres et al., 1987) or functions having both concave upwards and concave downwards regions (Clancy, 1992). In fact, an upwards curvature occurs when the considered nutrient limits herbivore growth. More generally, the curvature of the performance function depends on the considered nutrient, the range in which this nutrient is changed (Miles et al., 1982; Ohmart et al., 1985), and the age of the herbivore (Scriber and Slansky Jr, 1981; Ohmart et al., 1985; Montgomery, 1982). Nevertheless, such counterexamples do not necessarily contradict the conclusion by Wetzel et al. (2016), as long as the average overall performance of the herbivore population decreases with plant trait variability.

The straightforward conclusion by Wetzel et al. (2016) and Ruel and Ayres (1999) becomes less convincing when herbivore preference is taken into account. We found that herbivore populations benefit from high nutrient level variability irrespective of the curvature of the performance function when having considerable preference for leaves on which their performance is high. The intuitive explanation for these findings is that a herbivore having considerable preference can obtain more leaves with higher nutrient level when variability is larger, leading to a fitness increase as long as the cost for preference is not too high. Indeed, this type of herbivore preference is often observed (Via, 1986; Herrera, 2009; Tabashnik et al., 1981; Travers-Martin and Müller, 2008; Despres et al., 2007; Rausher, 1979; Leyva et al., 2003; Lubchenco, 1978; Mody et al., 2007). This means that a plant that is attacked by herbivores with a concave downwards performance function may adapt its strategy according to the strength of herbivore preference. However, when an adaptation includes a considerable cost and herbivore preference is close to the transition point (where the fitness changes little with the plant strategy), it may not be worth to change the strategy.

The only exception of these findings is the case where the performance function has a local maximum that coincides with the mean nutrient level, cp. Section A.1 in the Appendix. In this case herbivores benefit from low nutrient level variability independently of their preference. Non monotonic performance functions occur when performance is evaluated not with respect to the overall nutritional value but with respect to the concentration of a particular nutrient that leads to negative effects in excess concentration (e.g. because of a resulting unbalanced diet). Herbivores might perform best on leaves with the mean nutrient level when the herbivore population is well adapted to the plant and its nutrient concentration range.

Note that we tested the robustness of our results under changes of the preference mean in order to mirror that herbivores may not perfectly discriminate the leaf traits. We found the qualitative same results as long as herbivores prefer high-quality leaves (i.e. where herbivore performance is high).

All this means that the relation between the curvature of the herbivore performance function and the effect of nutrient level variability on herbivore fitness is more complex than previously thought. It requires a closer look at herbivore preference, the effect of which depends on (i) the cost for finding appropriate leaves (s. Fig. 7 in the Appendix), (ii) the sign and magnitude of the curvature of the performance function in the relevant trait range (s. Fig. 6 in the Appendix), and (iii) the plant nutrient level variability.

Indeed, empirical studies support these dependencies: They find that more specialized species have a stronger preference for leaves on which their performance is high than more generalized species (Gripenberg et al., 2010; Tilmon, 2008; Soto et al., 2012). In our model, herbivores that can grow well only on a relatively small nutrient level interval (because the performance function is concave upwards, or because its curvature is large) are the ones with the larger optimum preference because they benefit most from preference. Some studies find furthermore that the preference to oviposition leaves where the offspring performs well is stronger when high-quality resources are rare (Tilmon, 2008), in agreement with our finding that optimal herbivore preference increases with decreasing plant nutrient level variability (i.e. increasing *S*) when the performance function is concave upwards or linear as long as variability is not too low (which would probably be unrealistic anyway). When the performance function is concave downwards, this trend occurs also, as long as the cost for preference is low enough (see Fig. 7). We further found that herbivore fitness decreases with increasing nutrient level variability when herbivores show optimal preference (cp. Fig. 4). Such an optimal preference can develop when changes in the plant strategy occur slowly compared to adaptations of herbivore preference.

Our results do not necessarily imply that plant trait variability is a disadvantage for the plant when herbivores can exploit this variability by showing preference for high-nutrient leaves. Plant trait variability may benefit the plant population for reasons independent of the herbivore. For example, Kotowska et al. (2010) found that plant genetic diversity increases both plant and herbivore survival and biomass. Furthermore, the increase in plant biomass and survival was found in both the presence and absence of herbivory, whereby the percentage increase was lower in the presence of the herbivore (Kotowska et al., 2010). Hence, genetic diversity is beneficial for a plant population for other reasons than herbivory. Due to non-additive effects the productivity in genetic mixtures is not predictable by the productivity of the corresponding monocultures (i.e. each consisting of a genotype used in the genetic mixtures) (Kotowska et al., 2010). For instance, different resource uptake strategies may decrease intraspecific competition in genetic mixtures (Kotowska et al., 2010; Crutsinger et al., 2006).

Additionally, the plant population may benefit from large intraspecific plant trait variability in spite of herbivore preference when different herbivores have preference for different associated traits for instance due to differing specialization strategies (generalist vs. specialist) (Gutbrodt et al., 2012) or as a response to drought and associated changes in secondary defense compounds (Gutbrodt et al., 2011). In this case the mean preference of these herbivores is low.

Finally, we want to return again to the hypothesis by Wetzel et al. (2016) that the large plant nutrient level variability found in nature is *per se* beneficial for the plant because it leads to decreased herbivory due to Jensen’s inequality, as herbivore performance functions typically have a concave downwards curvature. In our study, we showed that this is true when the herbivore population has low preference for high-nutrient leaves, whereby the extent of preference depends on the three conditions listed above. We thus conclude that the hypothesis formulated in (Wetzel et al., 2016) may be true when herbivore preference has a low impact, e.g. when cost for preference is high or the curvature of herbivore performance function is small and/or negative. However, when herbivore preference is strong, nutrient level variability may not *per se* lead to decreased herbivory. As discussed above, this, however, does not necessarily imply that large nutrient level variability is an unfavorable evolutionary strategy for the plant.

To conclude, our study revealed the importance of considering trait variability in plants and that herbivore preference can considerably alter the impact of trait variability in a plant-herbivore system. Note that these results are also valid when considering other traits than the nutrient level of the leaves, as well as for intra- and inter-individual trait variability and for oviposition and feeding preferences in the larval stage. Our investigation did not include changes in the plant populations but considered the plant properties as being given. Further work needs to be done to analyze the coevolutionary outcome considering plant nutrient level variability and herbivore preference as strategies, for instance, by using evolutionary game theory (Maynard Smith, 1982; Drossel, 2001). Due to the complex relationships between plant trait variability, herbivore performance, and herbivore preference (potentially of different herbivores, that for instance differ in their specialization degree), the evolutionary outcomes will depend in a nontrivial way on the interplay of these features. Furthermore, competition between herbivore individuals for egg-laying sites (and/or food) becomes important when the available leaf density is limited, for instance because herbivore preference is strong. This will reduce the benefit of herbivore preference (Wetzel and Thaler, 2018), as the nutrients of highquality leaves must then be shared between several individuals. The present study represents an important foundation for such subsequent investigations.

## A Robustness tests

### A.1 Influence of a non monotonically increasing performance function

In empirical studies, the nutrient quality is usually not based on the overall nutritional value for the herbivore, but on the concentration of a specific nutrient such as nitrogen or phosphorous. Hence, empirical studies often find herbivore performance functions that have an optimum at an intermediate nutrient level (cp. Fig. 5(a), (c)) (Fischer and Fiedler, 2000; Joern and Behmer, 1997, 1998; Zehnder and Hunter, 2009), since an increase in this specific nutrient concentration may be associated with a lack of other nutrients and thus an unbalanced diet, or with a change in other chemical or physical properties (Tao et al., 2014). Hence, we test whether our results of Section 3.2 also apply for this kind of performance function. This means that the nutrient level *n* describes the concentration of a specific nutrient in this section.

**Fig. 5.**
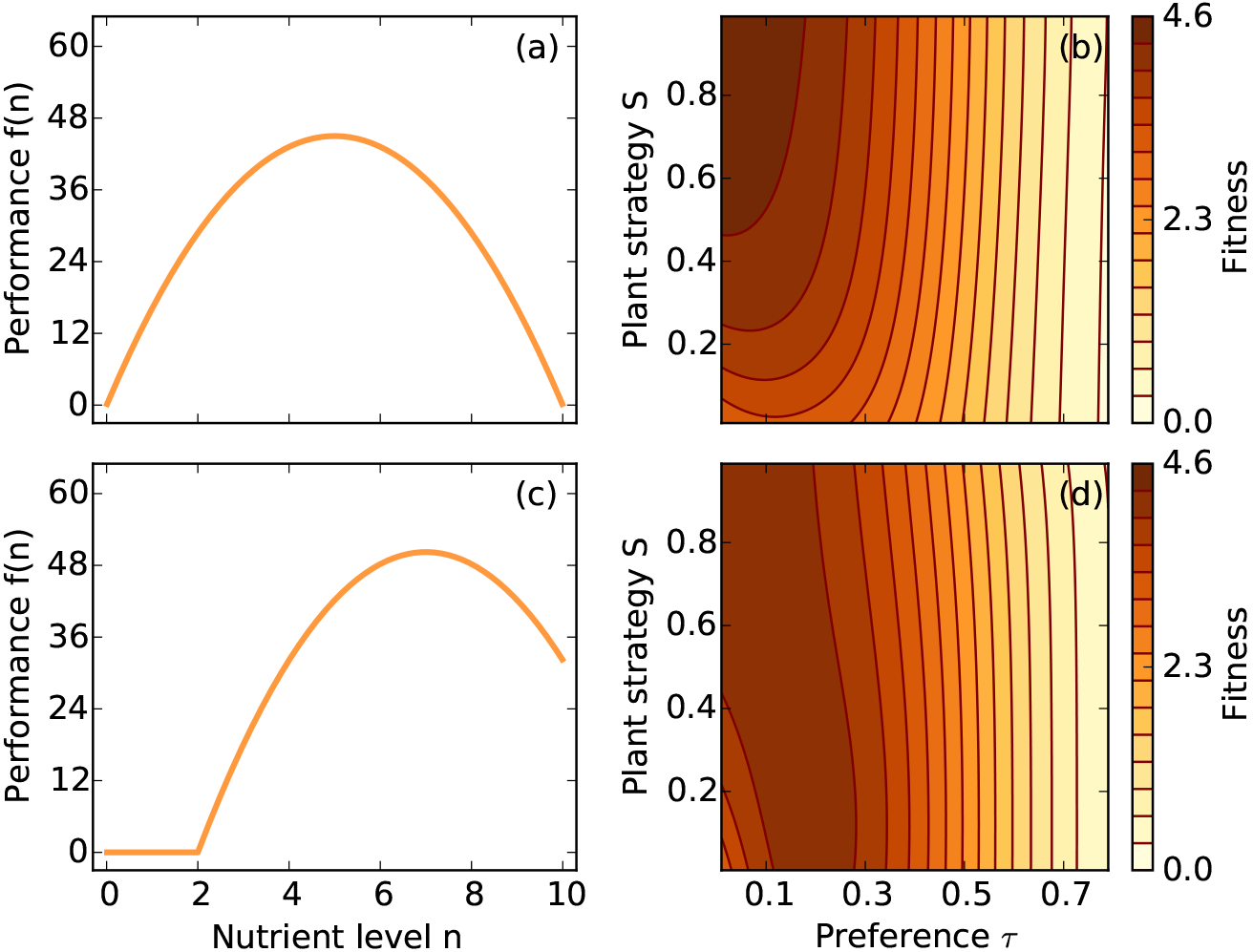
*(a)*,*(c)* The considered performance functions having its maxim um at an intermediate nutrient level: *(a)* 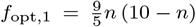, *(c)* 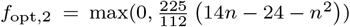. *(b)*,*(d)* Herbivore fitness (i.e. mean number of offspring per herbivore individual reaching reproductive age; cp. Eq.(7)) displayed in color code in dependence of the plant strategy parameter *S* (low *S* means high nutrient level variability, high *S* low variability; cp. Eq.(4)) and herbivore preference *τ* (cp. Eq.(5)) considering *f*_opt,1_ in (b) and *f*_opt,2_ in (d) as performance function.

Figs. 5(b), (d) show the mean fitness of a herbivore population 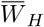 (cp. Eq.(1)) displayed in color code as a function of the herbivore preference *τ* (cp. Eq.(5)) and the plant strategy parameter *S* (cp. Eq.(4)) using the functions shown in Figs. 5(a), (c) as performance function, respectively. Note that the mean of the preference function again coincides with the maximum of the performance function and that we again normalized the mean performance of the two functions to 300 mg, i.e. 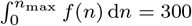, but that this normalization has no qualitative impact on the results.

The herbivore fitness decreases with increasing nutrient level variability (i.e. smaller *S*) when the nutrient level, where the performance function maximizes, coincides with the mean nutrient level 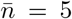 (s. Fig. 5(b)) as illustrated by the color change from darker to lighter color. This is because there are less leaves with the nutrient content preferred by the herbivore when nutrient distribution becomes broader. The figure also shows that a moderate degree of preference can increase the nutrient intake when the nutrient level variability is large (i.e. small *S*). When the mean nutrient level 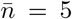 does not coincide with the maximum of the performance function (s. Fig. 5(c), (d)), herbivore populations having only a moderate degree of preference suffer from increased plant nutrient level variability (cp. Section 3.1). However, herbivore populations having a larger degree of preference (*τ* ⪆ 0.15) benefit from increased nutrient level variability (cp. Section 3.2). Hence, the results presented in Section 3.1 and 3.2 also apply to a performance function having its maximum at an intermediate nutrient level when this maximum does not coincide with the mean nutrient level 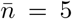.

### A.2 Influence of the magnitude of the curvature of the performance function

We showed in Section 3.2 and 3.2.2 that the sign of the curvature affects the extent of optimal herbivore preference. It is plausible that the magnitude of the curvature also has an effect on optimal herbivore preference. In order to test this hypothesis, we use the performance functions shown in Figs. 6(a), (c). The curvature is higher in (c) than in (a). A performance function with a larger curvature is for instance suitable when the herbivore is more specialized since it can only grow well on a smaller range of nutrient concentrations. Note that we again normalized the mean performance of the two functions to 300 mg, i.e., 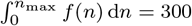, but that this normalization has no qualitative impact on the results.

**Fig. 6.**
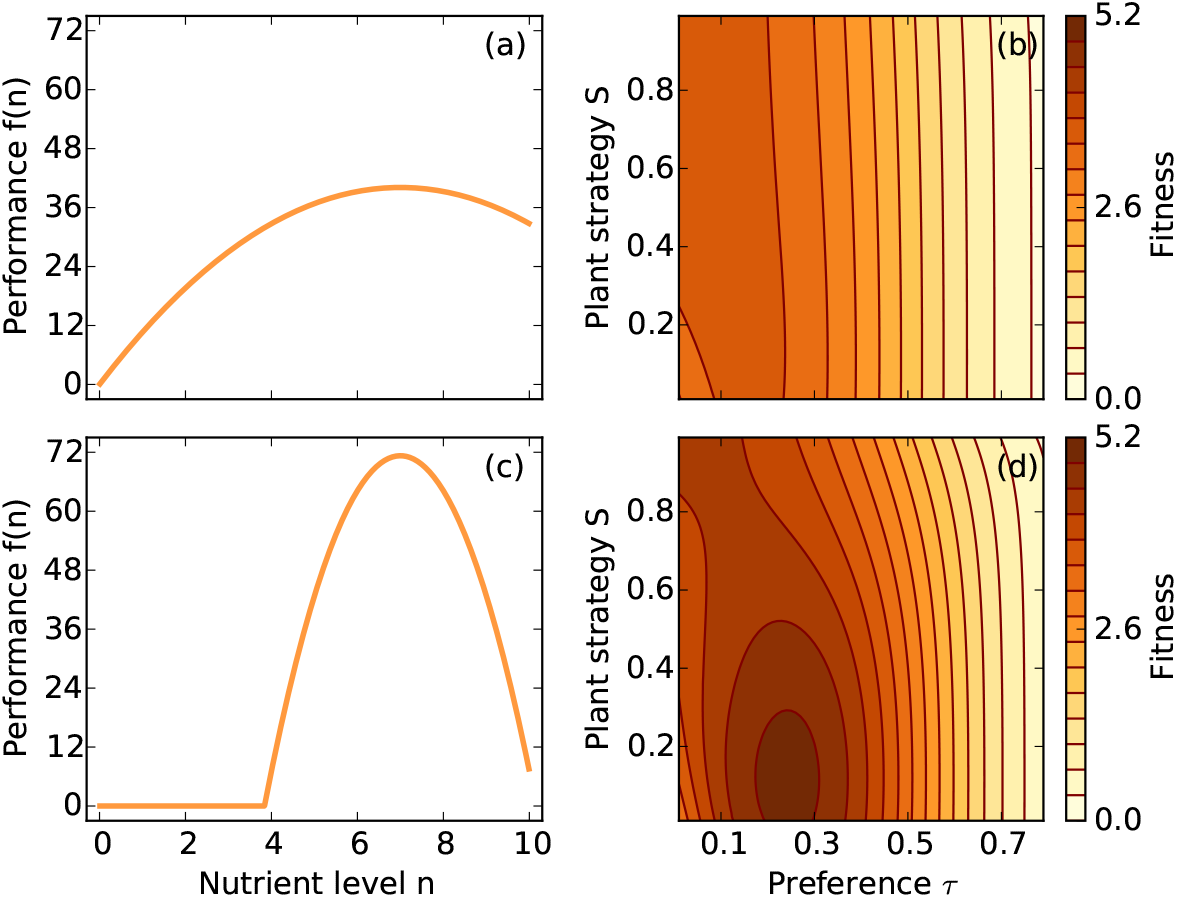
*(a)*,*(c)* The considered performance functions that differ in the magnitude of their curvature: *(a)* 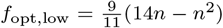, *(c)* 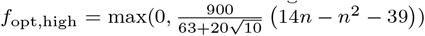 *(b)*,*(d)* Herbivore fitness (i.e. mean number of offspring per herbivore individual reaching reproductive age; cp. Eq.(7)) displayed in color code in dependence of the plant strategy parameter *S* (low *S* means high nutrient level variability, high *S* low variability; cp. Eq.(4)) and herbivore preference *τ* (cp. Eq.(5)) considering *f*_opt,low_ in (b) and *f*_opt,high_ in (d) as performance function.

Figs. 6(b), (d) show the mean fitness of a herbivore population 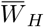 (cp. Eq.(1)) displayed in color code as a function of the herbivore preference *τ* (cp. Eq.(5)) and the plant strategy parameter *S* (cp. Eq.(4)), using these two performance functions.

The optimal herbivore preference is higher for all plant strategies *S* when the herbivore performance function has a higher curvature, i.e. *f*_opt,high_ (cp. 6(a), (c)) as can be seen by the location of the darkest color on the x-axis representing the preference *τ*. Furthermore, the fitness increase that can be achieved by having a preference is much larger when the curvature of the performance function is stronger. In this case, the herbivore benefits strongly from a broader nutrient distribution as there are much more leaves with the preferred nutrient content.

### A.3 Influence of the shape of the mass loss due to preference

The optimal herbivore preference for a given plant strategy parameter *S* (cp. Eq.(4)) depends on its costs. The shape of the mass loss due to preference *β*(*τ*) between the limits *τ* → 0 and *τ* → 1 is determined via the parameters *μ* and *k* (cp. Eq.(6)). Larger *μ* means that the cost of preference is larger, and large *k* means that the costs are mainly incurred when preference is large.

Fig. 7 shows the optimal herbivore preference *τ* (cp. Eq.(5)) for a given plant strategy parameter *S* (cp. Eq.(4)) for different values for *k* (left column) and for *μ* (right column) considering the concave upwards *f*_pos_(*n*) (diamonds), linear *f*_lin_(*n*) (circles), and concave downwards *f*_neg_(*n*) (triangles) performance function shown in Fig. 1(a). The color shades in the markers display the mean fitness of the herbivore population under the particular circumstances.

**Fig. 7.**
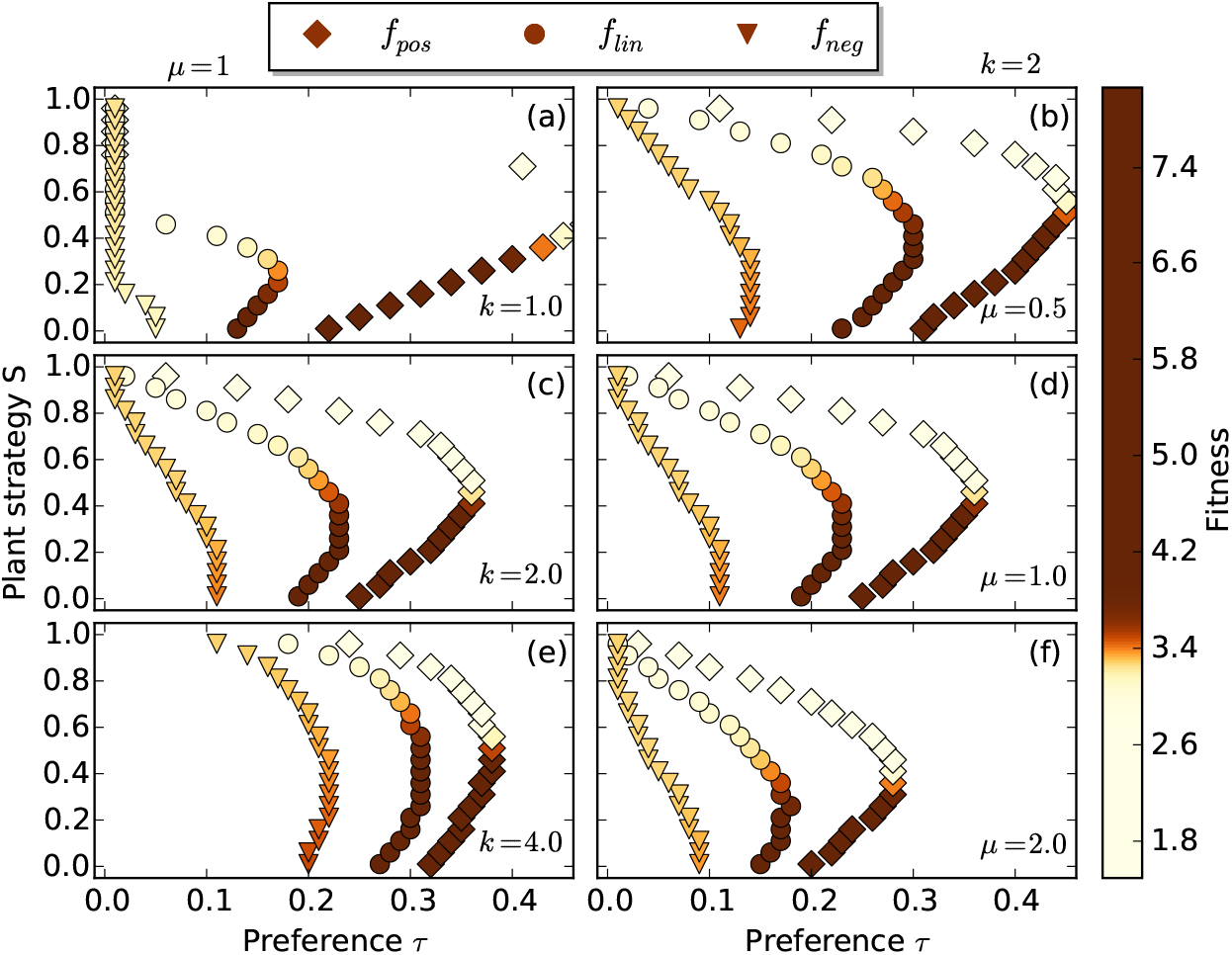
Herbivore preference *τ*, for which herbivore fitness is maximized for a given plant strategy parameter *S* (low *S* means high nutrient level variability, high *S* low variability; cp. Eq.(4)) considering different parameters *μ* ((a), (c), (e)) and *k* ((b), (d), (f)), that shape the mass loss due to preference *β*(*τ*) between the limits *τ* → 0 and *τ* → 1 (cp. Eq.(6)). Larger *μ* means that the cost of preference is larger, and large *k* means that the costs are mainly incurred when preference is large. The different markers represent the three different performance functions *f* (*n*) shown in Fig. 1(a). The color shades in the markers display the mean fitness of the herbivore population under the particular circumstances.

As in Section 3.2.2, we find that the optimal preference value *τ* is largest for the concave upwards performance function *f*_pos_(*n*), followed by the linear performance function *f*_lin_(*n*), and is smallest for the concave downwards performance function *f*_neg_(*n*) independent of the plant strategy parameter *S* (cp. Fig. 7).

The fitness, that is reached with optimal herbivore preference, increases with increasing *S* when *k* = 1 and herbivore performance is a concave downwards function *f*_neg_ (cp. Fig. 7(a)), since the herbivore population has very low or no preference in this case and thus benefits from low nutrient level variability (cp. Fig. 2(c)).

The larger *μ*, the lower is the optimal preference for a given plant strategy parameter *S* (cp. Fig. 7(b), (d), (f)) since the parameter *μ* determines where the mass loss *β*(*τ*) reaches its half saturation maximum (HSM). Half of the available mass will be lost to the cost of preference when

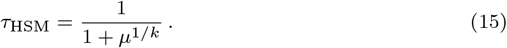

Hence, a larger value for *μ* implies a higher mass loss for a given preference *τ* (cp. Fig. 8(a)).

**Fig. 8.**
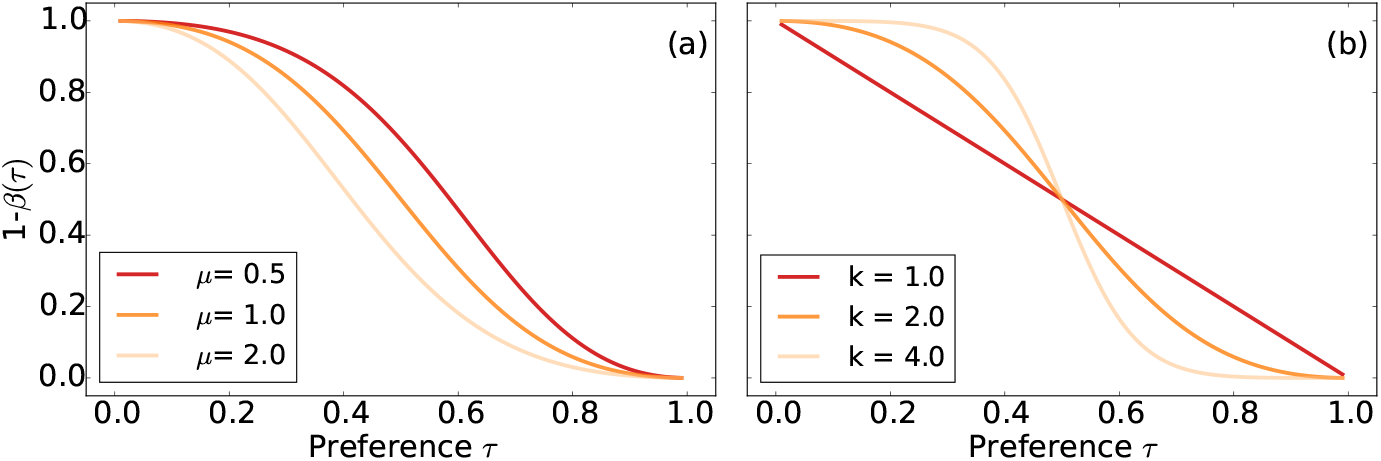
Proportion of mass that remains considering preference 1 − *β*(*τ*) for different values for *(a)* the parameters *μ* and *(b) k* that shape the mass loss due to preference between the limits *τ →* 0 and *τ →* 1. We chose *k* = 2 in (a) and *μ* = 1 in (b).

The exponent *k* determines the slope in the half saturation maximum (cp. Fig. 8(b)) and has a more divers impact on the optimal herbivore preference for a given plant strategy parameter *S*. More precisely, the optimal preference curves for different performance functions approach each other for increasing *k*. Under the assumption of a linear *f*_lin_ or a concave downwards *f*_neg_ performance function, the optimal herbivore preference decreases with decreasing *k* (cp. Fig. 7(d), (f)). In these cases, relatively low preferences *τ* lead to the highest fitness for a given plant strategy parameter *S* and in this preference range decreasing values for *k* lead to considerably higher losses due to preference (cp. Fig. 8(b)). The same is true in the case of a concave upwards performance function *f*_pos_ (cp. Fig. 7(b)) when *S* is small or high. For intermediate *S*, optimal preference is higher for *k* = 4 than for *k* = 2, since the resulting mass loss is smaller, however, optimal preference is highest for *k* = 1. In this range of *S*, optimal herbivore preference reaches higher values (near *τ* = 0.5), such that the relative mass loss *β*(*τ*) changes much faster with *τ* for increasing *k*. As a consequence, it pays off to have a stronger preference when *k* = 1. Nevertheless, one has to keep in mind that the maximal fitness values reached in this range for *S* are higher for larger values for *k*.

